# Mapping the human auditory cortex using spectrotemporal receptive fields generated with magnetoencephalography

**DOI:** 10.1101/2020.12.01.406843

**Authors:** Jean-Pierre R. Falet, Jonathan Côté, Veronica Tarka, Zaida-Escila Martinez-Moreno, Patrice Voss, Etienne de Villers-Sidani

**Affiliations:** Department of Neurology and Neurosurgery, McGill University

## Abstract

We present a novel method to map the functional organization of the human auditory cortex noninvasively using magnetoencephalography (MEG). More specifically, this method estimates via reverse correlation the spectrotemporal receptive fields (STRF) in response to a dense pure tone stimulus, from which important spectrotemporal characteristics of neuronal processing can be extracted and mapped back onto the cortex surface. We show that several neuronal populations can be found examining the spectrotemporal characteristics of their STRFs, and demonstrate how these can be used to generate tonotopic gradient maps. In doing so, we show that the spatial resolution of MEG is sufficient to reliably extract important information about the spatial organization of the auditory cortex, while enabling the analysis of complex temporal dynamics of auditory processing such as best temporal modulation rate and response latency given its excellent temporal resolution. Furthermore, because spectrotemporally dense auditory stimuli can be used with MEG, the time required to acquire the necessary data to generate tonotopic maps is significantly less for MEG than for other neuroimaging tools that acquire BOLD-like signals.

## Introduction

An important goal of auditory neurophysiology is to model the functional organization of the human auditory cortex (AC). This involves developing an intricate understanding of auditory processing along both spectral and temporal dimensions, and relating these features to the spatial topographical organization of the AC.

Frequently, the topographical organization of the human AC has been studied noninvasively using functional magnetic resonance imaging (fMRI) in terms of tonotopy, or best frequency maps, which has been found to be a key organizational feature (Da Costa et al., 2011; Formisano et al., 2003; Humphries, Liebenthal, & Binder, 2010; Langers & van Dijk, 2012; Talavage & Edmister, 2004; Woods et al., 2010). Although details such as the orientation of the tonotopic gradient are still debated, an anterior to posterior high-low-high best frequency organization centered on Heschl’s gyrus (HG) is found and agreed upon in most human fMRI studies (Gardumi, Ivanov, Havlicek, Formisano, & Uludağ, 2017), and is consistent whether pure tones or natural sounds are used (Moerel, Martino, & Formisano, 2012). Coupled with the spatial organization of other neuronal response characteristics such as the broadness of frequency tuning, and paired with findings from cyto- and myeloarchitectural studies, the AC has been further divided by fMRI into subfields with unique processing properties (Moerel, De Martino, Formisano, 2014).

However, the role of temporal processing within the micro-organization of the human AC remains unclear from the available fMRI literature alone (Leaver and Rauschecker, 2016). Crucial aspects of our sensory experience, such as speech perception and music enjoyment, clearly rely heavily on precise temporal encoding of auditory information (Abrams et al., 2011). Invasive electrophysiological recordings in several animal species have shown the importance of temporal features in understanding the functionality of AC subfields (Linden et al., 2003; Nagel & Doupe, 2008). Moreover, studying the temporal domain of auditory processing is necessary to gain a complete understanding of auditory plasticity (Schreiner & Polley, 2014; Carlin & Elhilali, 2015). For example, auditory training using temporal discrimination tasks can lead to improvements in the processing of temporal features that do not result in improvements in spectral processing (van Wassenhove & Nagarajan, 2007), reinforcing the importance of studying both dimensions. Similarly, studying temporal dynamics can yield insights into age-related changes in auditory processing (de Villers-Sidani et al., 2010).

Unfortunately, while fMRI boasts an excellent spatial resolution to answer questions pertaining to the spatial organization of the AC, it cannot provide sufficient temporal resolution to adequately study temporal dynamics and short-latency events. The hemodynamic response to neuronal activity measured by fMRI occurs on the order of seconds (Aguirre, Zarahn, & D’esposito, 1998), which precludes precise characterization of neuronal activity occurring on the order of milliseconds. Furthermore, because of the relatively long acquisition time, stimuli sets are typically small and offer less flexibility than one would ideally want to study the response to complex sounds. Studying auditory processing in fMRI has also been limited by loud operating noise, even though workarounds have been developed (Cha, Zatorre, & Schönwiesner, 2016).

MEG is an attractive alternative modality for *in vivo* electrophysiological recording of neuronal activity in the AC. It not only provides superior temporal resolution on the order of milliseconds (Regan, 1989), but also provides a completely silent acquisition environment. An important barrier preventing its widespread use has been related to concerns regarding its ability to spatially resolve the millimetric spatial organization of the AC (Langers & van Dijk, 2012; Moerel, De Martino, Formisano, 2014), in particular its tonotopic organization. This concern is offset by recent successes in capturing the retinotopic organization of the visual cortex using MEG at a spatial resolution of 7 mm in smooth cortical regions and less than 1 mm near curved gyri (Nasiotis, Clavagnier, Baillet, & Pack, 2017). Moreover, early efforts at identifying a basic tonotopic gradient using MEG have been successful in some respects. Dipole depth beneath the scalp has consistently been found to correlate with stimulus frequency, and orientation of the gradient has been shown to vary with gyral morphology (Romani, Williamson, & Kaufman, 1982; Pantev et al., 1988; Kuriki & Murase, 1989; Huotilainen et al., 1995; Verkindt, Bertrand, Perrin, Echallier, & Pernier, 1995). Other studies have also identified a posterior to anterior gradient, lower frequencies being represented more posteriorly, with the possibility of there being multiple tonotopic gradients (Pantev et al., 1995; Weisz, Wienbruch, Hoffmeister, & Elbert, 2004). Finally, a recent MEG study using speech sounds was able to identify a tonotopic gradient similar to that obtained in fMRI (Su, Zulfiqar, Jamshed, Fonteneau, & Marslen-Wilson, 2014).

Encouragingly, relatively simple study design tweaks could potentially yield improvements in the spatial resolution of MEG, notably through the use of higher stimulus density. There is evidence from research with owl monkeys pointing to an inverse relationship between stimulus density and the tuning width of neurons in the AC, as shown by the smaller size of their receptive fields with such stimuli (Blake & Merzenich, 2002). This could be due to increased peri-neuronal inhibition when stimuli are presented at a faster rate, increasing the spectrotemporal specificity of each neuron, and therefore improving the spatial resolvability of neuronal subpopulations. Using a dense stimulus could therefore improve the spatial resolution of MEG with respect to tonotopic organization.

Here, we describe a novel method to map the functional organization of the AC using MEG. Specifically, we take advantage of the MEG’s high temporal resolution to extract the spectral and temporal characteristics of sound processing for each neuronal source by computing their spectrotemporal receptive field (STRF), and demonstrate how the characteristics of STRFs can then be extracted and mapped onto the cortical surface to study organizational features such as tonotopy. STRFs have indeed been commonly used to describe the dynamics of neuronal activity in response to auditory stimuli (see for e.g.: Calabrese, Schumacher, Schneider, Paninski, & Woolley, 2011; Depireux, Simon, Klein, & Shamma, 2001; Kowalski, Depireux, & Shamma, 1996; Linden, Liu, Sahani, Schreiner, & Merzenich, 2003; Miller, Escabí, Read, & Schreiner, 2002; Nagel & Doupe, 2008; Sen, Theunissen, & Doupe, 2001; Theunissen, Sen, & Doupe, 2000; Woolley, Fremouw, Hsu, & Theunissen, 2005; Woolley, Gill, & Theunissen, 2006). They represent the spectral and temporal patterns of auditory stimuli that elicit the maximal response from a neuron. To estimate STRFs, several methods have been used for varying stimulus types (Theunissen, Sen & Doupe, 2000), but the foundational technique revolves around reverse correlation and involves averaging the stimulus content preceding neuronal spikes (de Boer & Kuyper, 1968). Doing so results in a spike-triggered average that can reliably estimate the STRF when using a stimulus that is uncorrelated in the spectral and temporal dimensions, as is typical for stimuli used for mapping tonotopy.

We show here that spectrotemporally dense auditory stimuli composed of isointensity pure tones (IIPTs) can yield sufficient spatial resolution to allow for mapping the tonotopic organization of the AC using reverse correlation-based STRFs generated from MEG. This method can therefore be reliably used to investigate the spatial organization of the AC, with the added benefit of MEG’s excellent temporal resolution to study short-latency-dependent events and complex spectrotemporal characteristics, permitting an in-depth non-invasive functional study of auditory processing in humans.

## Results

### I. Estimation of STRFs

For this analysis, we recorded the neural responses to a 10-minute IIPT stimulus train non-invasively in ten subjects (labeled S1 to S10) using a 275-channel whole-head MEG system (CTF MEG International Services Ltd.). MEG records magnetic fields outside the head, and a reverse problem must be solved to localize the source of the magnetic fields from where they originate inside the brain as electrical currents produced by neuronal activity. To do so, we used Weighted Minimum Norm Estimates (wMNE) (Lin et al., 2006) which constrains each source to a one-dimensional perpendicular orientation with respect to a cortex surface obtained through an MRI-based cortical reconstruction generated with FreeSurfer (Dale & Sereno, 1993). Our analysis was conducted on a high cortical tessellation (150,000 sources) to maximize the potential for high spatial resolution.

We generated STRFs for each source within our region of interest (ROI) in the right and left hemispheres of ten subjects (S1 to S10) using a technique based on reverse correlation analysis adapted to MEG data and detailed in *Materials and Methods*. The STRF represents the average stimulus-triggered activation amplitude (the average z-score value of every significant neuronal activation event). The resulting STRFs clearly display several important spectrotemporal characteristics expected of neurons in the AC (Figure 1). These include temporal features such as best temporal modulation rate and response latency, as well as spectral features such as best frequency and frequency bandwidth. The STRFs provide information about the auditory stimuli most likely to elicit a significant response from a given source. While the majority of STRFs had a single peak at a latency of about 100 ms (representing the M100 response), we could identify a number of sources that exhibited a smaller peak at a latency of 50 ms (representing the earlier M50 response). Some sources exhibited complex STRF spectrotemporal patterns, including some with frequency sweeps.

**Figure 1.**
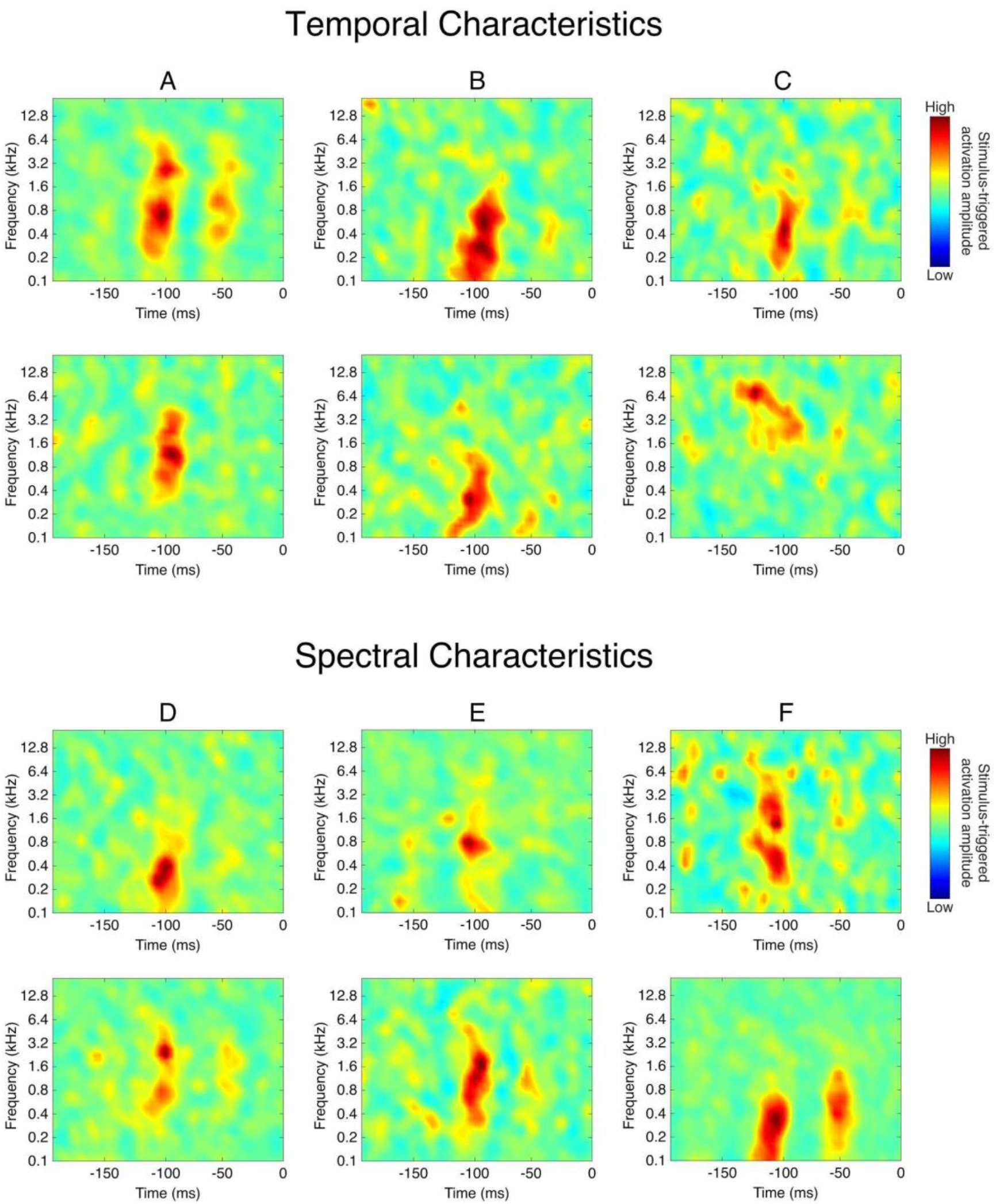
Variety of STRF characteristics. Sample STRFs exhibiting a range of spectral and temporal characteristics. (A) M50 response (top = strong response; bottom = no response). (B) Best temporal modulation rate (top = low modulation rate; bottom = high modulation rate). (C) Temporal complexity (top = preference for a temporally isolated stimulus; bottom = preference for a downward frequency sweep). (D) Best frequency (top = low frequency; bottom = high frequency). (E) Frequency bandwidth (top = small bandwidth; bottom = large bandwidth). (F) Spectral complexity (top = two spectral peaks eliciting an M100 response; bottom = two spectral peaks eliciting an M100 and/or an M50 response).

Key properties that can be obtained through analysis of STRFs are shown in Figure 2. These histograms represent a group-level average among all subjects. Best frequencies were represented along a bimodal distribution with one peak at 0.283 kHz and another at 0.8 kHz. However, the range was large, extending throughout all presented frequencies. On average, 90% of sources per subject had a best frequency between 0.2 and 3.2 kHz. Frequency bandwidths were most commonly 2.5 octaves, with the remainder of sources exhibiting a large range of bandwidth. M100 latency was most commonly at 110 ms. Finally, best temporal modulation rate also followed a bimodal distribution, with one peak at 15 Hz (with rates ranging from 10 to 24 Hz), and another centered around 33 Hz (with rates ranging from 25 to 100 Hz).

**Figure 2.**
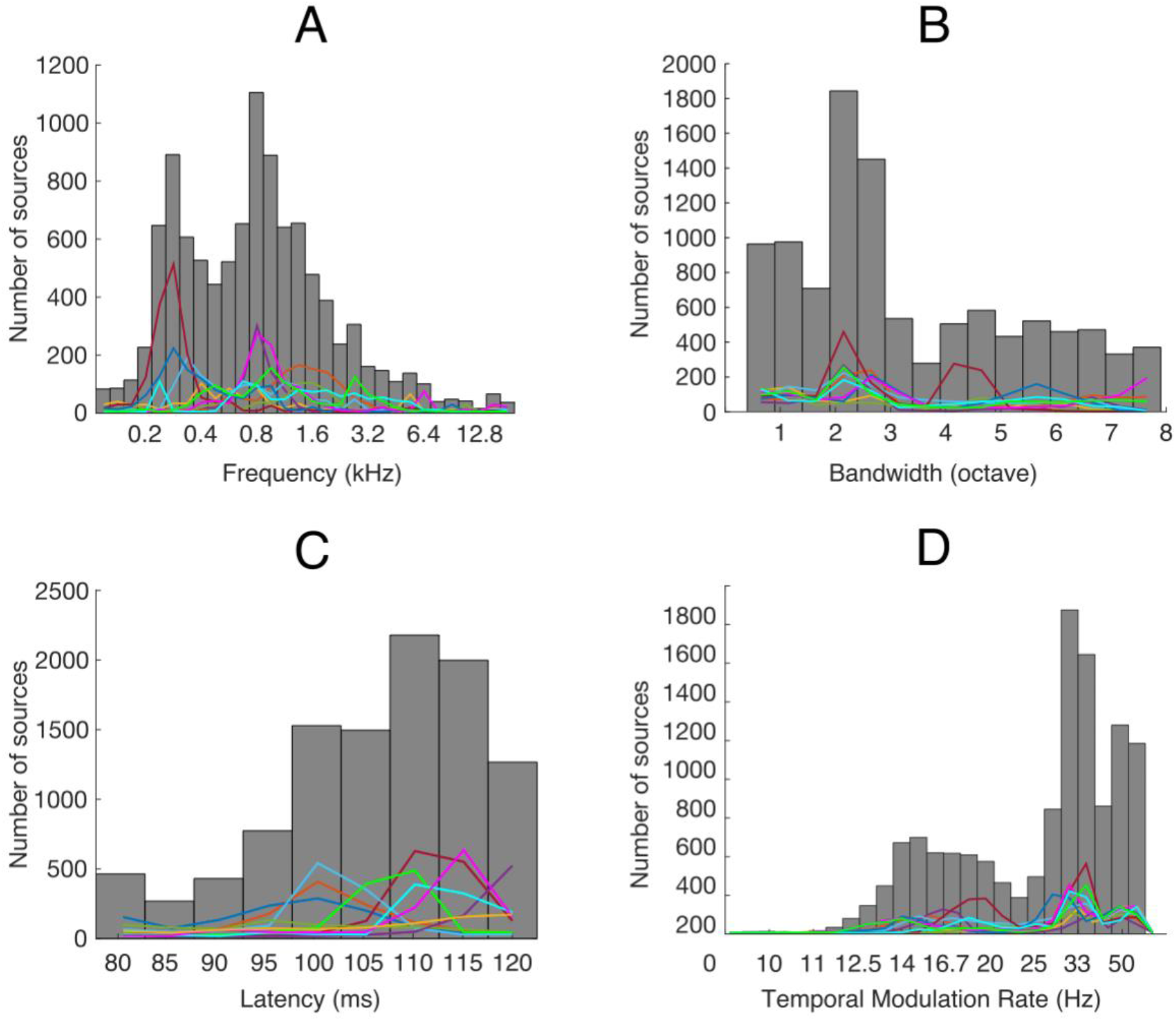
Histograms of STRF characteristics. Histograms showing the total number of sources from all 10 subject for each of the following STRF characteristics: best frequency (A), bandwidth (B), latency (C), and best temporal modulation rate (D). Lines representing the individual contribution of each subject are superimposed onto the histograms.

### II. Selection of IIPT-Responsive Sources

We defined IIPT-responsive sources as those having an STRF M100 response peak greater than a z-score of 3.5, with a latency between 80-120 ms, and a minimum STRF bandwidth of 0.375 octaves (see *Materials and Methods* for precise definitions). The high z-score threshold enables the selection of only those sources that are very IIPT-responsive. This threshold is should be determined based on the amount of smoothing that is used in the STRF-generation and the signal-to-noise ratio of an experiment. The latency thresholds enable the identification of the M100 response with a range of response latencies. Finally, the bandwidth threshold enables the selection of physiologically plausible receptive fields, eliminating sources that could have a significant “single-bin” receptive field due to chance alone, given the high number of data bins present in the STRF.

### III. Identification of a tonotopic gradient

To demonstrate the utility of computing STRFs in MEG to study the spatial topographic organization of the auditory cortex, we generated tonotopic maps from the best frequency values of the STRFs for each IIPT-responsive neuronal source. A tonotopic organization could be identified in the right temporal lobe for all subjects, as shown in Figure 3. Because of variability between subjects in the position of tonotopic gradient reversals and in the underlying cortical anatomy which covers only a very small area, we do not show a group-level average using currently available tools in the Brainstorm suite, as this leads to loss of valuable gradient information. The gradient pattern is best analyzed individually or, alternatively, using a manual landmark-based averaging method which has proven successful in some fMRI studies (e.g. Humphries et al. 2010).

**Figure 3.**
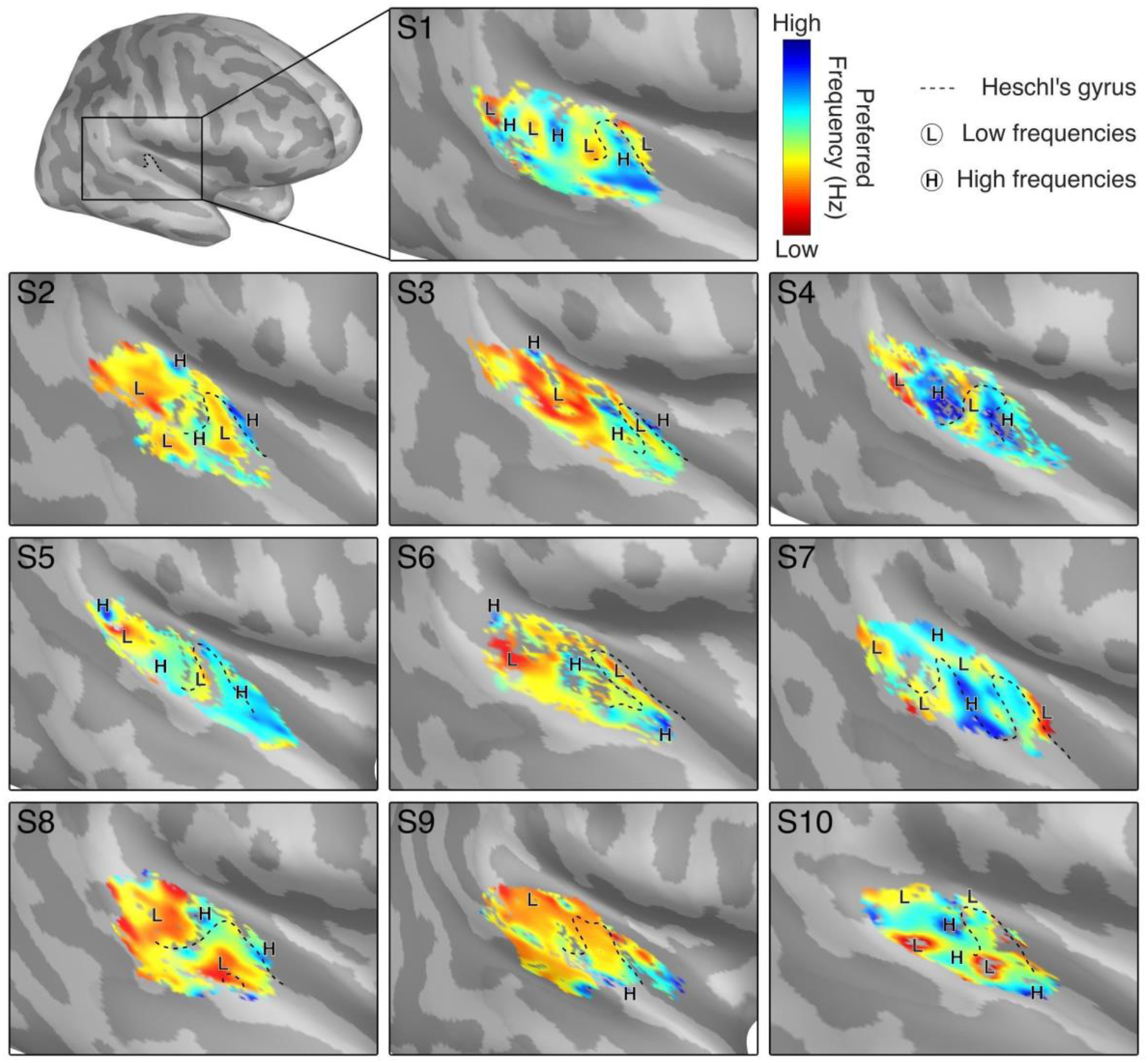
Best frequency maps and tonotopic gradient organization. Best frequency maps are shown for subjects 1 to 10. Major regions of high and low frequencies are marked as H and L, respectively, and Heschl’s gyrus is outlined for reference. Tonotopic gradients can be identified in all subjects. Colormap limits are set near the local minima and maxima of each subject to best visualize gradient patterns.

For the majority of subjects (S1 to S8), a primary tonotopic gradient perpendicular to the longitudinal axis of HG could be identified. This primary gradient is most often centered on the posterior part of HG. Among the two subjects who did not have a perpendicular gradient progression, S9 had a simple antero-posterior gradient oriented parallel to the longitudinal axis of HG, and S7 had several circular zones of low and high frequencies with a complex organization not observed in other subjects. Of note, all subjects had a single HG, while S7 had a complete duplication of HG, and S8 had a partial duplication of HG. There was a more variable tonotopic organization present in planum temporale (PT), usually with a relative overrepresentation of low frequencies.

Other characteristics of STRFs can be projected onto the cortical surface, including bandwidth, latency, and temporal modulation. The right-hemisphere maps for these characteristics are presented in Figure S1, S2, and S3.

### IV. Investigation of lateralization to IIPT stimuli

Identification of tonotopic gradients was more robust in the right cerebral hemisphere of subjects, which is why the remainder of our analysis was performed on the right. Best frequency maps for the left hemisphere are shown in Figure S4.

The left hemisphere’s tonotopic maps had a decreased signal-to-noise ratio, a shorter range of best frequencies, and less elaborate tonotopic gradients with some subjects having no discernible gradient. To investigate whether this was associated with a difference in the number of IIPT-responsive sources in each hemisphere, we calculated the percentage of IIPT-responsive sources within each hemisphere’s ROI (Figure 4). While there were over 50% of IIPT-responsive sources in the left hemisphere’s ROI, a two-tailed paired t-test revealed that the right hemisphere had 24.3% more IIPT-responsive sources than the left hemisphere (95% CI: 14.8 - 33.8; p = 0.0003), confirming a lateralization to the right.

**Figure 4.**
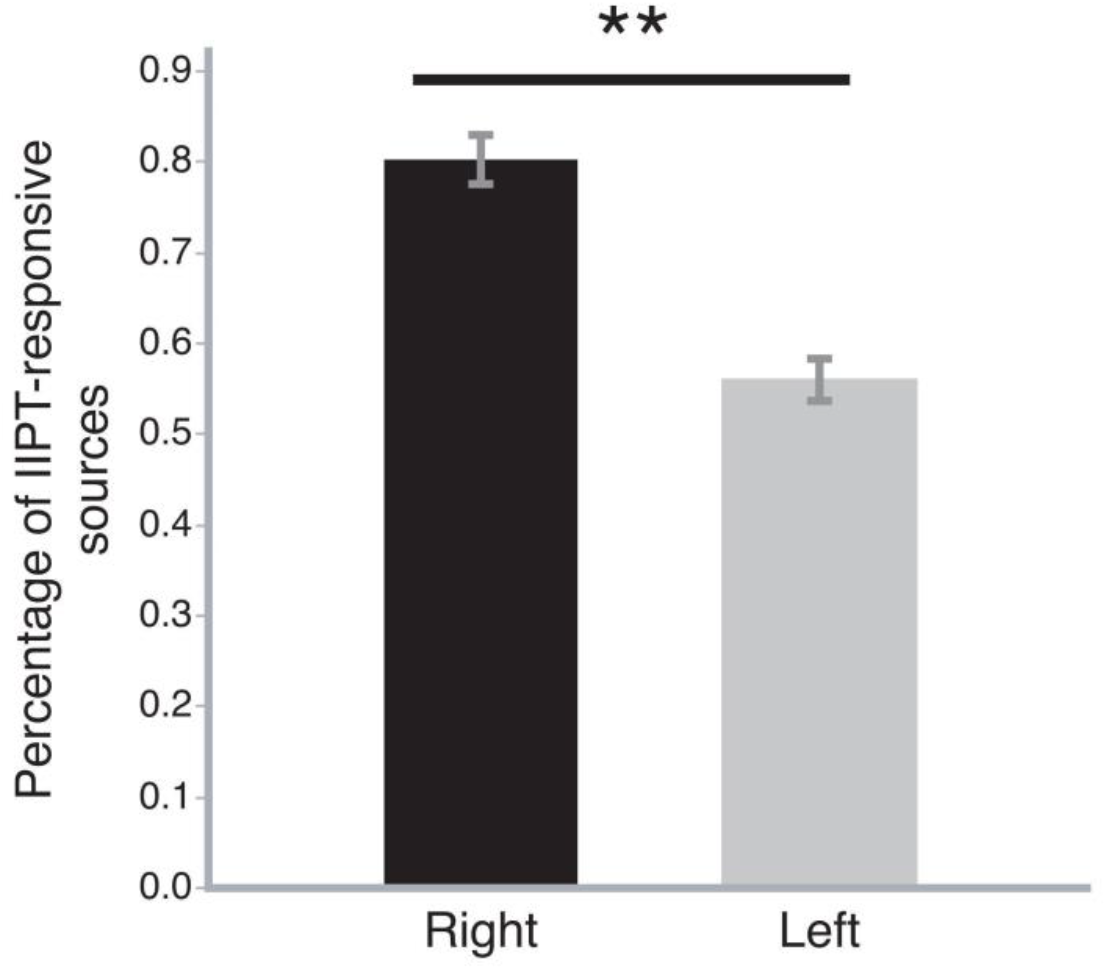
Average percentage of IIPT-responsive sources for each cerebral hemisphere among all subjects. The average percentage of IIPT-responsive sources is presented for the right and left hemispheres. The percentage represents the number of IIPT-responsive sources over all possible sources within our region of interest. The error bars represent the standard error to the mean. Two-tailed paired t-test results in p = 0.0003, t-value −5.7887, DF = 9.

## Discussion

We have described a novel method for functional mapping of the human AC using MEG, showing that it can reliably extract important information about the spatial organization of auditory processing when using STRFs generated from a dense pure tone auditory stimulus. This supports the hypothesis that the spatial resolution of MEG is more than sufficient to study tonotopic gradients in the human AC, and allows us to leverage the MEG’s excellent temporal resolution to study short-latency-dependent events and complex spectrotemporal characteristics inherent in auditory processing that are impossible to study using fMRI.

### I. Estimation of STRFs using MEG

MEG has been used in the past to successfully generate physiologically plausible STRFs using discrete (Constantino et al., 2017) and continuous stimuli (Crosse et al., 2016; Ding & Simon, 2012), but to the best of our knowledge, this is the first *in vivo* MEG study to estimate STRFs using reverse correlation in human subjects for the purpose of mapping the functional organization of the AC. Our findings support that it is possible to generate physiologically plausible STRFs with excellent variety in terms of spectrotemporal patterns, akin to what has been reported in other mammalian studies (see for e.g.: Elhilali, Fritz, Chi, & Shamma, 2007; Massoudi, Van Wanrooij, Versnel, & Van Opstal, 2015). Some STRFs exhibited complex patterns, both spectrally and temporally, which is a testament to the high temporal resolution and sufficient spatial resolution of MEG to be able to isolate such a large variety of neuronal sub-populations in a relatively small cortical region. In some cases, we could even detect a strong M50 response.

Our analysis of the distribution of these STRF properties revealed a bimodal distribution of best frequency centered on 0.283 kHz and 0.8 kHz, with the vast majority of sources (90%) having a best frequency between 0.2 kHz and 3.2 kHz. Our results are comparable with those of a small study of four epilepsy patients with intracranial electrode recordings, where only the frequencies 0.32 kHz to 3.2 kHz elicited neuronal responses, and 0.25 to 2.0 kHz elicited the strongest responses (Bitterman, Mukamel, Malach, Fried, & Nelken, 2008). This finding likely reflects emphasis on frequencies used in speech sounds. In an articulation test (intelligibility of speech communication), subjects scored 95% accuracy when a low-pass filter of 4.0 kHz was applied to speech sounds and 100% when the low-pass filter was 7.0 kHz (Monson, Hunter, Lotto, & Story, 2014), suggesting that spectral content below 7.0 KHz is the most important for speech comprehension. The wider audible frequency range in humans extends up to about 20 kHz for young healthy individuals (Monson et al., 2014), and while these were represented in our dataset, they were markedly underemphasized when compared to the frequency band of speech. One possible confounder pertains to the E-A-RTONE 3A insert earphones, which have frequency responses that, although audible, progressively decreases in intensity beyond 3 kHz. Although this could contribute to the underrepresentation of higher frequencies in our dataset, the fact that frequency representation begins dropping well below 3.0 kHz (starting at 0.8 kHz) suggests that this finding is truly representative of the underlying functional organization.

The ability to extract STRFs using MEG is significant. The STRFs we have produced are in keeping with what is physiologically expected of neurons in the human AC. The spectrotemporal characteristics of auditory processing are likely important in understanding the subdivisions of neuronal populations in humans. STRF analyses have also been crucial to better understand the mechanisms underlying plasticity using the AC as a model, particularly in mice. In this setting, STRFs have provided an excellent means of visualizing the changes in spectrotemporal characteristics of neuronal response over time (Kamal, Holman, & de Villers-Sidani, 2013). Until now, correlating STRFs with spatial organization in the auditory cortex was only possible in animal studies and intracranial recording studies in humans, but our proposed methodology provides a novel non-invasive method to do so in humans.

### II. MEG-generated tonotopic maps

A mirror-symmetric tonotopic gradient has been described in most fMRI studies (Da Costa et al., 2011; Formisano et al., 2003; Humphries et al., 2010; Langers & van Dijk, 2012; Moerel, De Martino, Formisano, 2012), and the majority of our subjects exhibit a very similar gradient pattern, usually centered around a region of low frequency in the posterior part of HG. The directionality of the gradient found using our method is most closely aligned with the findings of Humphries et al., Da Costa et al., Formisiano et al., and Moerel et al. (Humphries et al., 2010, Da Costa et al., 2011, Formisano et al., 2003; Moerel, De Martino, Formisano, 2012), who also describe a primary gradient perpendicular to the longitudinal axis of HG (though Langers & van Dijk (2012) in contrast describe a latero-medial progression). Furthermore, a review publication integrating fMRI research with cyto- and myeloarchitectural studies proposed a model of the human AC with a tonotopic gradient oriented at a similar angle with respect to HG (Moerel, De Martino, Formisano, 2014). Therefore, the fact that the tonotopic organization we describe is in keeping with what is found in the fMRI literature supports the accuracy of the findings generated by our technique. It adds to the body of evidence pointing to a primary gradient that is perpendicularly oriented to the longitudinal axis of HG, which likely represents the primary AC. This primary gradient is most often centered on HG.

There was significant intersubject variability in our dataset, which is consistent with the findings of methods boasting greater spatial resolution such as fMRI (Humphries et al., 2010). Nonetheless, we could still identify consistent tonotopic gradient progressions that shared similar patterns and directionality among the majority of subjects. These patterns extend from the core auditory cortex to the putative location of the belt and parabelt areas. These other subfields have been characterized by relying on spatial organization of best frequency (Moerel, De Martino, Formisano, 2012). The technique presented here has the spatial resolution that would allow further characterization of these subfields’ features by harnessing the MEG’s temporal resolution.

### III. Right-hemispheric lateralization of response to pure tones

The tonotopic maps produced using our methodology have led us to identify a right-hemispheric lateralization of the tonotopic organization in response to IIPTs at M100. Despite there being over 50% of sources in the left hemisphere that responded to the IIPT stimulus, the characterization of a tonotopic organization was less robust than in the right hemisphere, with some subjects having no discernible gradient. While functional lateralization of the human AC has been extensively studied with respect to stimuli involving music and speech sounds (Tervaniemi & Hugdahl, 2003), lateralization of tonotopy using pure tone stimuli has received less attention in the literature. In a single fMRI study, the presence of a clearer tonotopic organization was noted in the right primary AC compared to the left, although there was significant inter-subject variability (Langers, Backes, & van Dijk, 2007). Right-sided specialization for frequency-specific tuning has also been noted in intracranial recordings of auditory evoked potentials (Liégeois-Chauvel, Giraud, Badier, Marquis, & Chauvel, 2001) and in a previous study using MEG (Ozaki & Hashimoto, 2007). There is also evidence pointing to left-ear advantage (and therefore right hemispheric lateralization) when human subjects are presented with tonal, but not noise stimuli (Sininger & Bhatara, 2012).

The bulk of the evidence from the available literature studying pitch and music points to the right hemisphere having better spectral resolution (Zatorre, Belin, & Penhune, 2002), and therefore implicating it more in music, pitch and tonal processing. This contrasts with the left hemisphere’s better temporal resolution, rendering it more important in the processing of much faster temporal variations in the sound amplitude envelope, as is the case in speech. These hypotheses are supported by lesioning studies showing that lesions affecting the right HG result in deficits in the perception of pitch, by electrophysiological studies showing an association between pitch perception and the timing of cortical activity in the right hemisphere, and by a variety of functional imaging studies showing a predilection for tonal processing in the right hemisphere (Zatorre, Evans, & Meyer, 1994; Zatorre, Evans, Meyer, & Gjedde, 1992; Perry et al., 1999; Halpern & Zatorre, 1999; Griffiths, Johnsrude, Dean, & Green, 1999; Penhune, Zattore, & Evans, 1998; Hugdahl et al., 1999; Tervaniemi et al., 2000).

Several fMRI studies have identified a tonotopic organization in the left hemisphere (see for e.g. Formisano et al., 2003; Talavage et al., 2004; Langers et al., 2007). While we could identify a tonotopic organization in a subset of participants’ left hemispheres, this was less robust than on the right. This discrepancy could be due to at least two reasons. First, it is possible that the type of stimulus could be implicated. We used a spectrotemporally dense pure tone stimulus with a much greater rate of stimulus presentation than is typically used in fMRI studies. However, because the left hemisphere is thought to be important in the processing of temporal characteristics of sound (Zatorre et al., 2002), it would be difficult to explain why such a difference in the stimulus presentation rate could result in a lateralization to the right hemisphere. Second, the discrepancy could be related to the timing of acquisition and the temporal resolution of the two modalities. With MEG, the high temporal resolution allows us to isolate specific auditory cortical responses such as the M100 response, whereas the BOLD response used in fMRI results from neuronal activity occurring over a much longer time period, dictated by hemodynamic properties. Therefore, the activity captured through fMRI may relate to activity taking place much later than the M100 response in the auditory processing hierarchy. We believe this to be the more likely explanation behind this observation.

### IV. Limitations

There are limitations to the method we propose. First, the sound intensity (volume) of stimulus presentation is a limiting factor in the ability to resolve a tonotopic gradient. In order to truly capture the characteristic (best) frequency of a neuron, the lowest sound intensity that will elicit a response must be found; however, current electrophysiological and functional neuroimaging techniques are not sensitive enough to record neuronal responses barely above threshold, and therefore require the use of higher sound intensities. Coupled with the notion that neurons respond to a broader range of frequencies when stimulated by higher sound intensities (Recanzone, 2000), doing so may result in the spread of activation limiting the accuracy and resolvability of the measured tonotopic gradient (Tanji et al., 2010). While this limitation cannot be avoided, we used A-weighted stimulus intensity to compensate for the differences in volume necessary to lead to equivalent intensity perception at each frequency (Fletcher & Munson, 1933).

There are possible artifacts related to recording auditory evoked fields in the region of the AC. MEG is selectively sensitive to current along the walls of sulci, and cannot detect current at the crest of gyri and bottom of sulci (Puce & Hämäläinen, 2017). Moreover, the activity recorded from regions lying in close proximity to other surfaces, as is the case with the AC, could potentially be altered or even canceled by conflicting currents occurring simultaneously on the adjacent surface (Ahlfors et al., 2010). In our dataset, we did not observe any deficiency in the identification of IIPT-responsive sources in the crests of gyri and bottom of sulci. This leads us to believe that any potential alteration in signal occurring as a consequence of the macroanatomy of the AC did not prevent adequate source estimation with MEG.

Although our analysis is based on the earliest consistently detectable response in MEG (Pantev et al., 1988), the M100 response, what it represents remains controversial. Intracranial recordings have localized M100 to the lateral portion of HG and PT (Godey, Schwartz, de Graaf, Chauvel, & Liégeois-Chauvel, 2001; Liégeois-Chauvel, Musolino, Badier, Marquis, & Chauvel, 1994), while non-invasive recordings have localized it exclusively to PT (Lütkenhöner & Steinsträter, 1998; Engelien, Schulz, Ross, Arolt, & Pantev, 2000), which may be interpreted as activity in secondary ACs. This evidence rightfully has led some to question the claim that M100 originates from the primary AC (Moerel, De Martino, & Formisano, 2014). However, our data is not entirely consistent with this view. We show that there is clear activity at M100 along the purported anatomical location of the primary AC, HG. There are two possible ways to reconcile these differences. It may be that the spatial resolution of MEG is such that activity in spatially separated cortical areas appears to be overlapping. In this case, most of the observed activity could be originating from PT but falsely appear to be extending beyond PT into HG. We believe this is unlikely, particularly given that tonotopic gradients were identified as progressing in shorter distance increments than the distance between PT and HG. Another possibility is that primary and secondary auditory processing are overlapping in some regions of the AC. If this were the case, it would indicate that HG is both involved in primary and secondary processing. There is evidence showing that the earlier 50 ms-latency response (M50) is in fact located within the same anatomical region as the M100 response (Wang et al., 2014), which could support this hypothesis. Even if M100 represents higher order processing, we assume, as others have (Su et al., 2014), that the tonotopic organization of auditory processing should in theory remain stable over at least several hundred milliseconds. Even if it does not, investigation of the M100 response using MEG remains valuable, as insights into later auditory processing steps can be gained from studying the response in secondary ACs.

### V. Conclusions

Here, we show that MEG can be used to characterize the tonotopic organization of the AC by generating STRFs with a spectrotemporally dense pure tone stimulus. We described a large variety of STRF patterns consistent with the expected variety of neuronal subtypes that can be further studied both spectrally, through measures such as frequency bandwidth, and temporally, through measures such as best temporal modulation rate, and latency. The best frequency maps and tonotopic gradients we were able to generate shared strong similarities with those observed in other fMRI studies. MEG therefore is able to provide sufficient spatial resolution to study the spatial functional organization of the human AC, including the microarchitecture of auditory subfields, while providing additional benefits through its high temporal resolution. Our proposed method has significant implications for the field of auditory processing, as it is the first to effectively capture both high spatial resolution and spectrotemporal information, which together provide a more complete understanding of auditory processing in humans.

## Materials and Methods

### I. Participants

Ten right-handed subjects were recruited into the study (henceforth labeled S1 to S10). Three were female and the average age was 23 (range 19-27). All subjects reported being free of hearing impairment or neurological conditions that could affect brain function, including mild cognitive impairment, dementia and previous history of stroke. All subjects provided written informed consent. This study was approved by the research ethics board of the Montreal Neurological Institute.

The MEG and anatomical MRI recordings of S3 are freely available for download from the OpenNeuro platform at the following link: https://openneuro.org/datasets/ds003082/versions/1.0.0

### II. MEG Analysis

#### II.a. Stimuli presentation

Stimuli were generated by a Sound Blaster X-Fi Titanium HD audio card (Creative, Jurong East, Singapore) connected to a pair of E-A-RTONE 3A insert earphones (3M company, Indianapolis, Indiana).

The stimulus train was a 10-minute train of 50-ms long gated IIPTs (Figure 5). Thirty two different frequencies were presented with A-weighted intensities for the resulting stimulus train to be perceived at a similar intensity (Fletcher & Munson, 1933). More specifically, A-weighting describes the decibel attenuation necessary for each frequency to be perceived at the same intensity, because the perceived intensity varies depending on the frequency of the stimulus. There was an average of 55 pure tones per frequency, per recording, totaling 1,795 pure tones per recording. Frequencies ranged from 0.1 kHz to 25.6kHz, each separated by a quarter of an octave. The inter-stimulus interval was randomly generated from a gamma distribution with shape parameter 6 to achieve an average presentation rate of 3 Hz. Tones could overlap but less than 1% of tones did, and only 2 tones for every 64 were adjacent.

**Figure 5:**
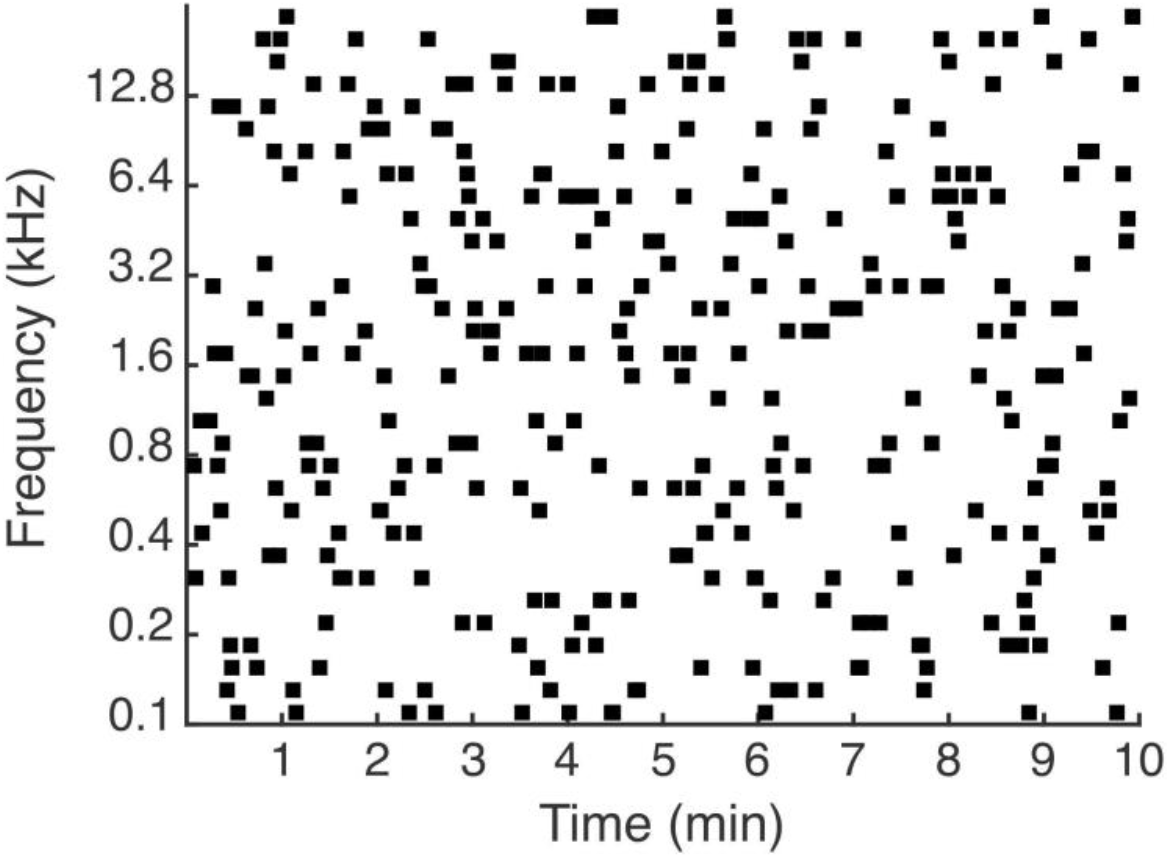
Iso-intensity pure tones stimulus. Sample stimulus spectrogram used to obtain STRFs. Frequencies range from 0.1 kHz to 25.6kHz and each is separated by a quarter of an octave. Tones are presented binaurally at an average rate of 3 Hz for a total of 10 minutes. Note that the width of the squares in this figure is larger than their true duration (50 ms) for the sake of presentation, and more tones appear to be overlapping than is truly the case. Refer to Figure 6. A for a true representation of the duration of each stimulus over a smaller time-window.

Subjects were instructed to fixate on a visual fixation cross throughout the stimulus presentation to reduce eye movement artifacts. The volume intensity was set to a comfortable hearing level.

#### II.b. MEG Acquisition

Using a 6 degrees-of-freedom digitizer (Patriot - Polhemus; Matlab interface RRID: SCR_006752) each subject's head was digitized. The head shapes contained about 100 to 200 points distributed across the scalp, eyebrows and nose to precisely coregister the activity to the structural MRI. Three coils were attached to fiducial anatomical locations on the head (nasion, and left and right pre-auricular points) to capture head movement inside the MEG. To record blinks and eye movements, we placed bipolar electro-oculographic (EOG) leads about 1 cm above and below one eye, and about 1 cm lateral of the outer canthi. Electrocardiographic (ECG) activity was recorded with one channel. The electrical reference was placed at the opposite clavicle. Both EOGs, ECG and the electrical reference were used for subsequent MEG artifact detection and removal. MEG was recorded using a 275-channel (axial gradiometers) whole-head MEG system (CTF MEG International Services Ltd.). All data were downsampled to 2400 Hz.

#### II.c. Structural MRI

Three-dimensional T1-weighted anatomical MR image volumes covering the entire brain were acquired on either a 1.5T Siemens Sonata or 3T Siemens Magnetom Prisma scanner with an 8 channel head coil (repetition time = 27 ms; echo time = 9.20 ms; between 176 and 192 sagittally oriented slices with slice thickness of 1 mm; acquisition matrix = 240×256; field of view = 256 mm).

#### II.d. MEG Data Pre-Processing and Spatial Modeling

MEG data analysis was performed in Matlab (RRID: SCR_001622; MATLAB and Statistics Toolbox Release 2015b), coupled with the Brainstorm extension (Tadel, Baillet, Mosher, Pantazis, & Leahy, 2011), which is documented and freely available for download online under the GNU general public license (RRID: SCR_001761; Tadel, 2019). MRI-based cortical reconstruction and volumetric segmentation were performed with the FreeSurfer image analysis suite (RRID: SCR_001847; Fischl, 2013; Dale & Sereno, 1993; Fischl, Sereno, & Dale, 1999; Fischl, Liu, & Dale, 2001).

Raw MEG data was pre-processed to remove signal contamination due to ocular, cardiac, and muscular artifacts using signal-space projections (Tesche et al., 1995; Uusitalo & Ilmoniemi, 1997). Each recording was then manually reviewed to discard any segment still experiencing significant contamination from artifacts.

The forward problem was solved using the overlapping-sphere approach (Huang, Mosher, & Leahy, 1999), which fits a sphere to the scalp surface. This simplified modeling method can be used given that the magnetic fields recorded from the brain are not distorted by the shape of the skull (Barth, Sutherling, Broffman, & Beatty, 1986; Okada, Lahteenmäki, & Xu, 1999). wMNE (Lin et al., 2006) was used to solve the reverse problem, with sources being constrained to a one-dimensional perpendicular orientation with respect to the cortex surface. The MRI-based cortex surface was generated with FreeSurfer and contained 330,000 sources (Dale & Sereno, 1993). Otherwise, default Brainstorm parameters were used in the wMNE modeling (SNR: 3 / Whitening: PCA; Regularize noise covariance: 0.1; Depth weighting: Order 0.5 / Maximal amount 10).

To reduce computation time, a lower resolution cortical tessellation (15,000 sources) was used to generate the wMNE source model for the purpose of regional time-frequency analysis. A high resolution cortical tessellation (150,000 sources) was used for the remainder of the analysis to maximize the spatial resolution.

#### II.e. Time-Frequency Decomposition

A time-frequency (TF) decomposition was done to select the optimal band-pass filter to apply to the pre-processed IIPT recording before further analysis. This analysis was conducted on all subjects using a randomly selected subset consisting of 10% of the presented IITPs. An anatomical ROI was selected for the TF decomposition. Given the putative primary AC’s location over HG (Liegeois-Chauvel, Musolino, & Chauvel, 1991), the ROI was based on the Desikan-Killiany parcellation for HG generated by FreeSurfer (Desikan et al., 2006), which was then manually enlarged to cover the surrounding sulcal space on both hemispheres. Using the 15,000 source-model, the analyzed ROI overlying HG covered an average of 320.6 sources (SD 23.7) or 49.6 cm^2^ (SD 4.28) per subject.

The recording was divided into trials of 1 s, from −500 to 500 ms with respect to each IIPT. The DC offset was corrected using the 500 ms period before each IIPT as a baseline. Time-series for each source within the ROI were extracted for each trial. These time-series were then subjected to a TF-decomposition using Morlet wavelets (Tallon-Baudry & Bertrand, 1999) characterized by a central frequency of 1Hz and a time resolution of 1 s. The decomposition was analyzed in 1 Hz-sized frequency bins. These parameters were chosen to maximize the spectral resolution at the 100 ms response latency (M100), with the goal of using the M100 response for the remainder of the analysis. The M100 is the earliest detectable event-related response attributable to the AC that can be reliable measured with auditory evoked fields (Pantev et al., 1988). Moreover, the M100 response measured by MEG correlates well spatially with the response measured through intracranial recordings (Godey et al., 2001).

The resulting TF-decompositions were then normalized by z-score transformation using a 250 ms-baseline before each IIPT, and an average across all ROI sources for each subject was obtained. A conservative z-score threshold of 1 was applied to the average TF-decomposition to identify the information-containing frequency bands at the 100 ms latency. Across all subjects, the minimum lower cutoff frequency was 3 Hz and the maximum upper cutoff frequency was 13 Hz (for the average TF-decompostition, see Figure S5; for the individual subjects’ TF-decomposition values, see Table S1). A band-pass filter of 3-13 Hz with a stopband attenuation of 60 dB was therefore applied to the pre-processed IIPT recordings for further analysis.

#### II.f. Estimation of STRFs

The ROI was constructed with the Desikan-Killiany parcellation of the transverse temporal gyrus (HG) and the part of the superior temporal gyrus that is posterior to HG, given that our focus was on the purported region of the primary auditory cortex. STRFs were generated using a technique based on the reverse correlation approach for each source within our ROI. Figure 6 depicts this process.

**Figure 6.**
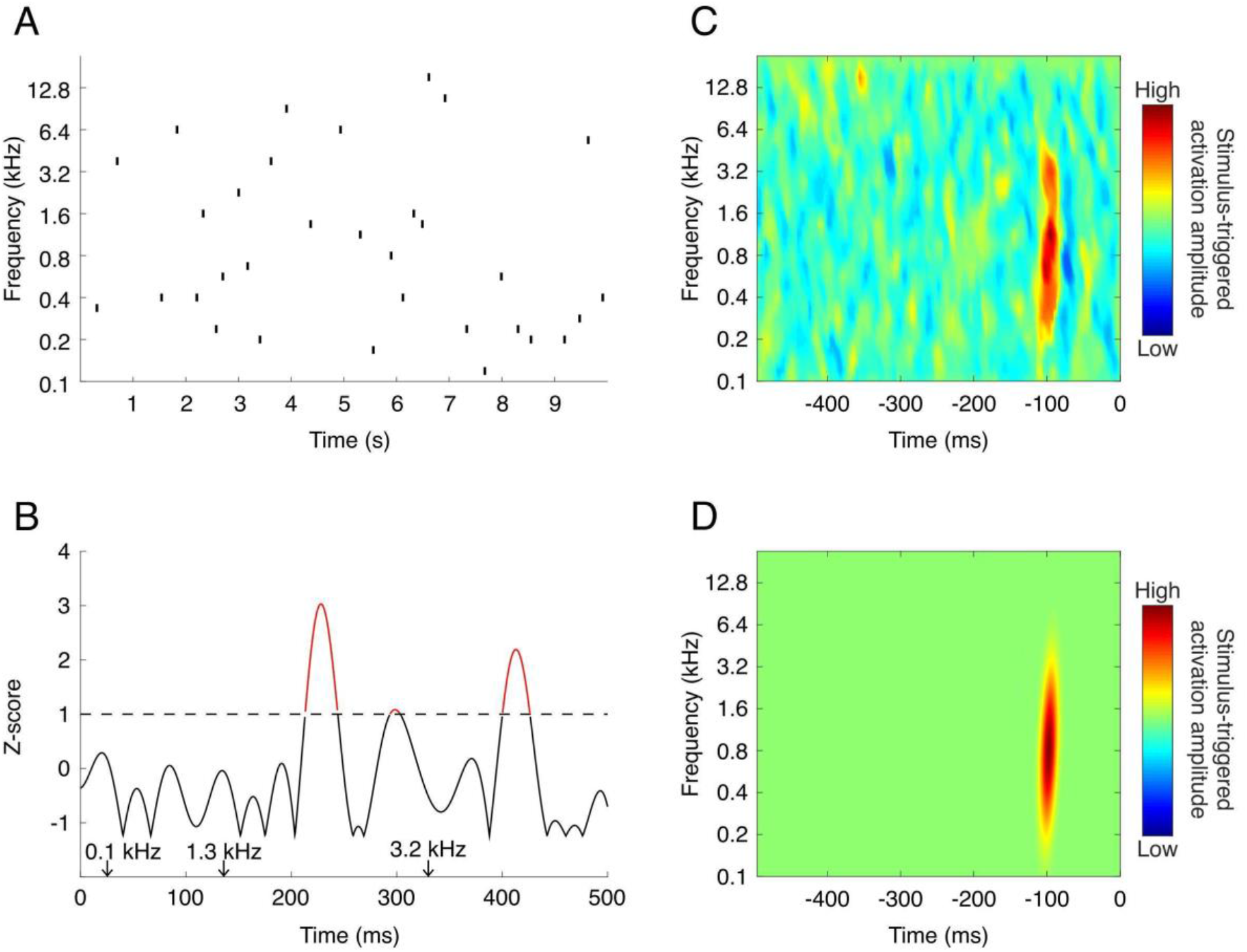
Generation of STRFs. The process of generating an STRF is depicted. (A) 10-second sample of the IIPT recording. (B) Z-score transformed source-space time series. The threshold for defining a significant activation event is shown with a dashed line at a z-score of 1. Significant activation events are shown in red. The time points at which three sample pure tones were played are shown with arrows above the x-axis. (C) Sample STRF, representing the average stimuli preceding all significant activation events for a given source. This particular STRF has a best frequency of 1.3 kHz. (D) Gaussian-fitted STRF.

The source-space time series was first extracted and converted to absolute values to eliminate the effect of the dipole current’s directionality which is not of interest for our application. The absolute value of the time series was then transformed into a z-score normalized time series dynamically by recalculating the mean and standard deviation at each reasonably long segment of silence. This segment of silence had to be at least 100 ms long and begin a minimum of 350 ms after the end of the previous IIPT stimulus to avoid contaminating the baseline with late stimulus-related responses.

From the z-score transformed source-space time-series, local maxima were extracted. A significant activation event was defined as a local-maximum with z-score > 1 (shown in red in Figure 6, panel B). Such a conservative z-score threshold was chosen to avoid missing activation events that could be reliably time-locked to a stimulus but that may have an amplitude that is relatively low. This choice is counterbalanced by the fact that we weigh activations proportionally to their z-score amplitude, as explained below. We believe this low z-score threshold in combination with a weighting system leads to a more objective selection of significant activations. In comparison, choosing a higher z-score threshold arbitrarily to select a smaller number of activations could be highly dependent on the signal-to-noise ratio of a particular experiment, where different thresholds may lead to different tonotopic maps.

To calculate STRFs, a method based on reverse correlation analysis (deCharms, Blake, & Merzenich, 1998; de Boer & Kuyper 1968) was used. Reverse correlation analysis can be used to reliably estimate a neuron’s STRF when the stimulus is uncorrelated, or sampled randomly and uniformly across the spectrotemporal dimensions as is the case with our IIPT stimulus (Theunissen, Sen & Doupe, 2000). In summary, the STRF produced through reverse correlation represents the linear estimate of the optimal stimulus preceding a neuronal activation event. It is calculated by computing the average stimulus, in both spectral and temporal dimensions, that precedes a neuronal activation event. For several authors (see for e.g. deCharms, Blake, & Merzenich, 1998), this neuronal activation event is a spike rate, and the STRF quantity is therefore a stimulus-triggered spike rate average. In the method described below, we used a stimulus-triggered activation amplitude average (the average z-score value of the significant activation events), which is more in keeping with the metric being recorded by MEG. The importance of a given activation event on the resulting STRF is therefore proportional to its amplitude.

More specifically, this STRF was computed as a matrix *STRF*(*f*, *t*), where *f* represents each 32 presented stimulus frequencies and *t* represents 4 ms bins within the 500 ms time window preceding a significant activation event. For each significant activation event *i*, the stimulus content in the preceding 500 ms time window was extracted. For each stimulus with frequency *f* and time *t* within this time-window, a value corresponding to the z-score amplitude of the corresponding significant activation event was defined as *Z_i_*(*f*, *t*). This z-score amplitude was then corrected for the slight variation in the total number of stimuli presented for each stimulus frequency by multiplying it by the coefficient 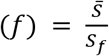, where 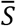 represents the mean number of stimuli presented per stimulus frequency, and *S_f_* represents the total number of stimuli presented of frequency *f*. The corrected z-score activation amplitudes corresponding to each stimulus within the reverse correlation time-windows were then summated to generate the final matrix representing the average stimulus-triggered activation amplitude:

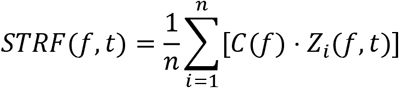

The final STRF was smoothed using a gaussian-weighted moving average with a window size of 4×4. The M100 response was then defined as the highest spike within a latency window of 80 to 120 ms. The STRF’s best frequency was defined as the frequency that elicited the maximal amplitude of activation at a latency corresponding to the M100 response. The best frequency was z-score transformed using the segment of the STRF from −500 to −350 ms for the purpose of determining which STRF showed a significant M100 response (see *Results, Selection of IIPT-Responsive Sources*).

The STRF was finally fitted to a 2D-gaussian surface, aligned on the peak corresponding to the M100 response, in order to smooth the data for estimation of bandwidth and best temporal modulation rate. To normalize its value according to the overall amplitude, the bandwidth was defined as the full spectral width at half maximum of the gaussian-fit and represents the range of frequencies that can elicit an M100 response. The best temporal modulation rate was calculated as *R* = (2*W*)^−1^, where *W* represents the temporal width of the gaussian-fit. The best temporal modulation rate represents a source’s preference for a stimulus with a particular temporal modulation.

Only sources with best frequencies ranging from 0.119 kHz to 18.102 kHz were included in the subsequent analysis (total of 30/32 frequencies). The frequency extremes were eliminated to eliminate the edge-effect caused by smoothing the STRFs.

The Brainstorm process used to generate STRFs and map the STRF features onto a cortex surface is available under an open source BSD license at the following GitHub repository: https://github.com/NeuroSensoryBiomarkingLab/MEGACmapping.

## Acknowledgements

We thank Sylvain Baillet, PhD, Robert Zatorre, PhD, and Kuwook Cha, PhD, from the Department of Neurology and Neurosurgery at McGill, for providing helpful comments about our methods and analysis.

## Additional Files

Supplementary File 1

