## Supplementary File 1 for "Mapping the human auditory cortex using spectrotemporal receptive fields generated with magnetoencephalography"

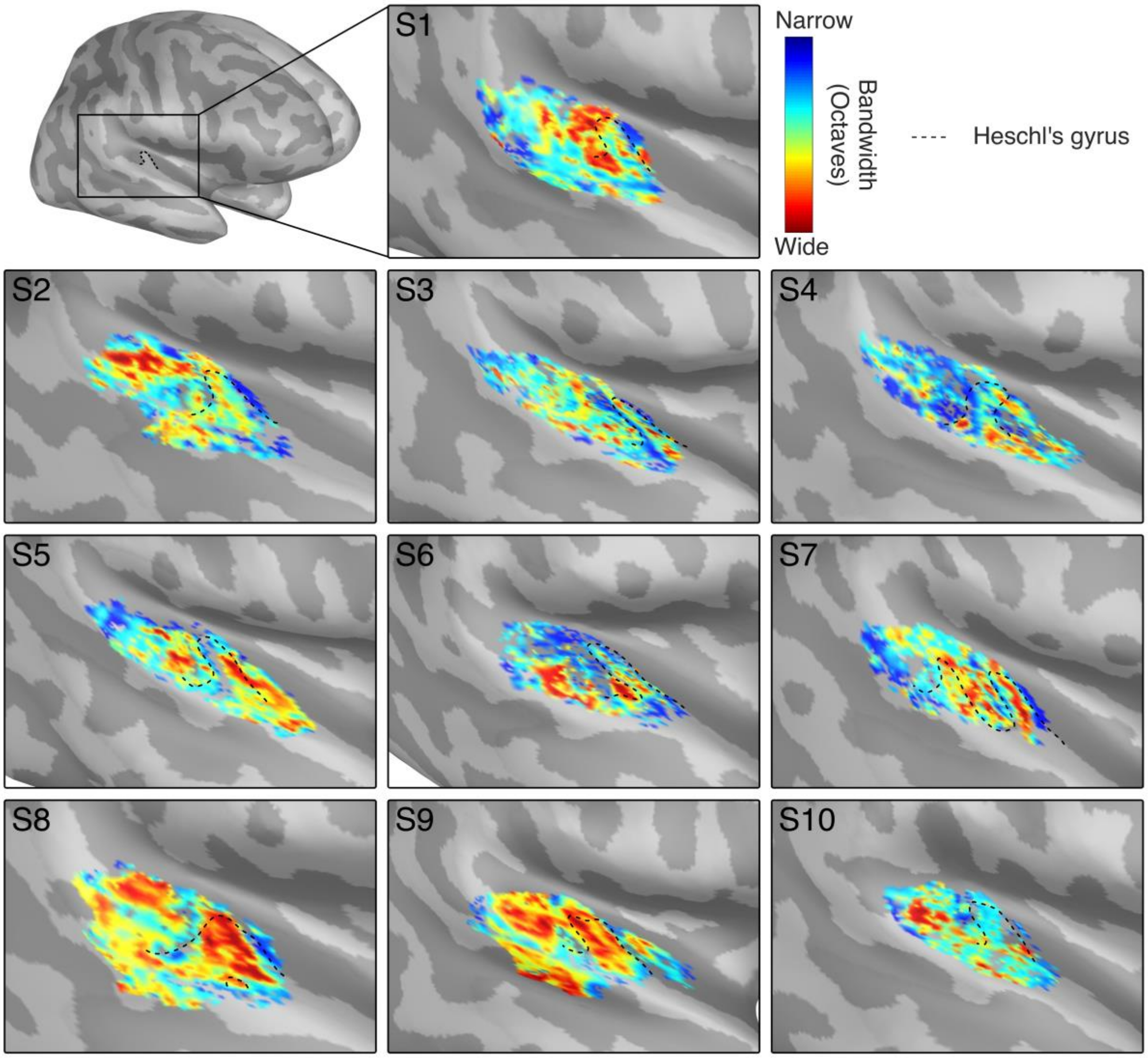


***Figure S1. Right Hemisphere Bandwidth Maps.*** *Bandwidth maps are shown for subjects 1 to 10. Heschl's gyrus is outlined for reference. Colormap limits are set near the local minima and maxima of each subject to best visualize gradient patterns.*

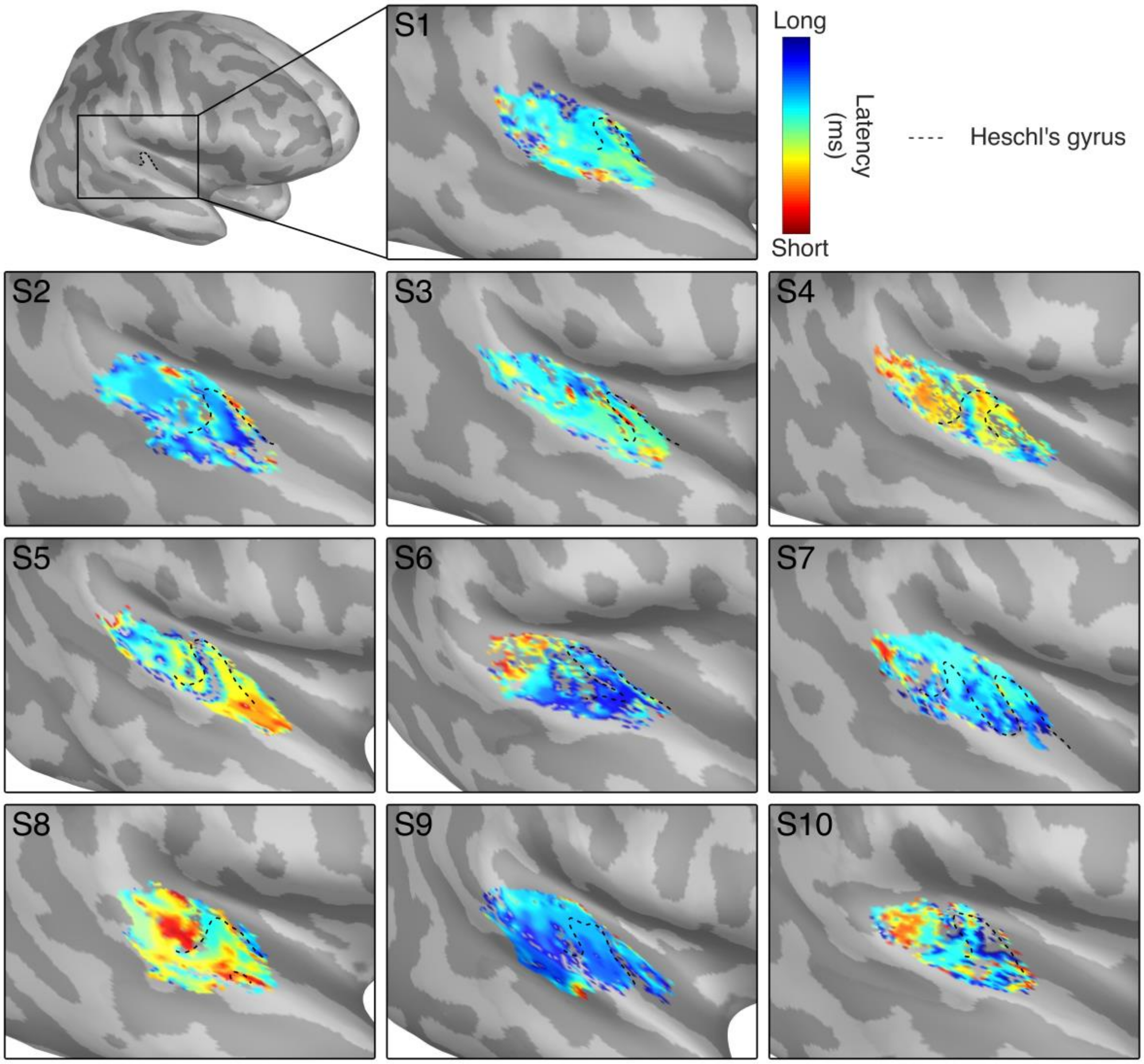


***Figure S2. Right Hemisphere Latency Maps.*** *Latency maps are shown for subjects 1 to 10. Heschl's gyrus is outlined for reference. Colormap limits are set near the local minima and maxima of each subject to best visualize gradient patterns.*

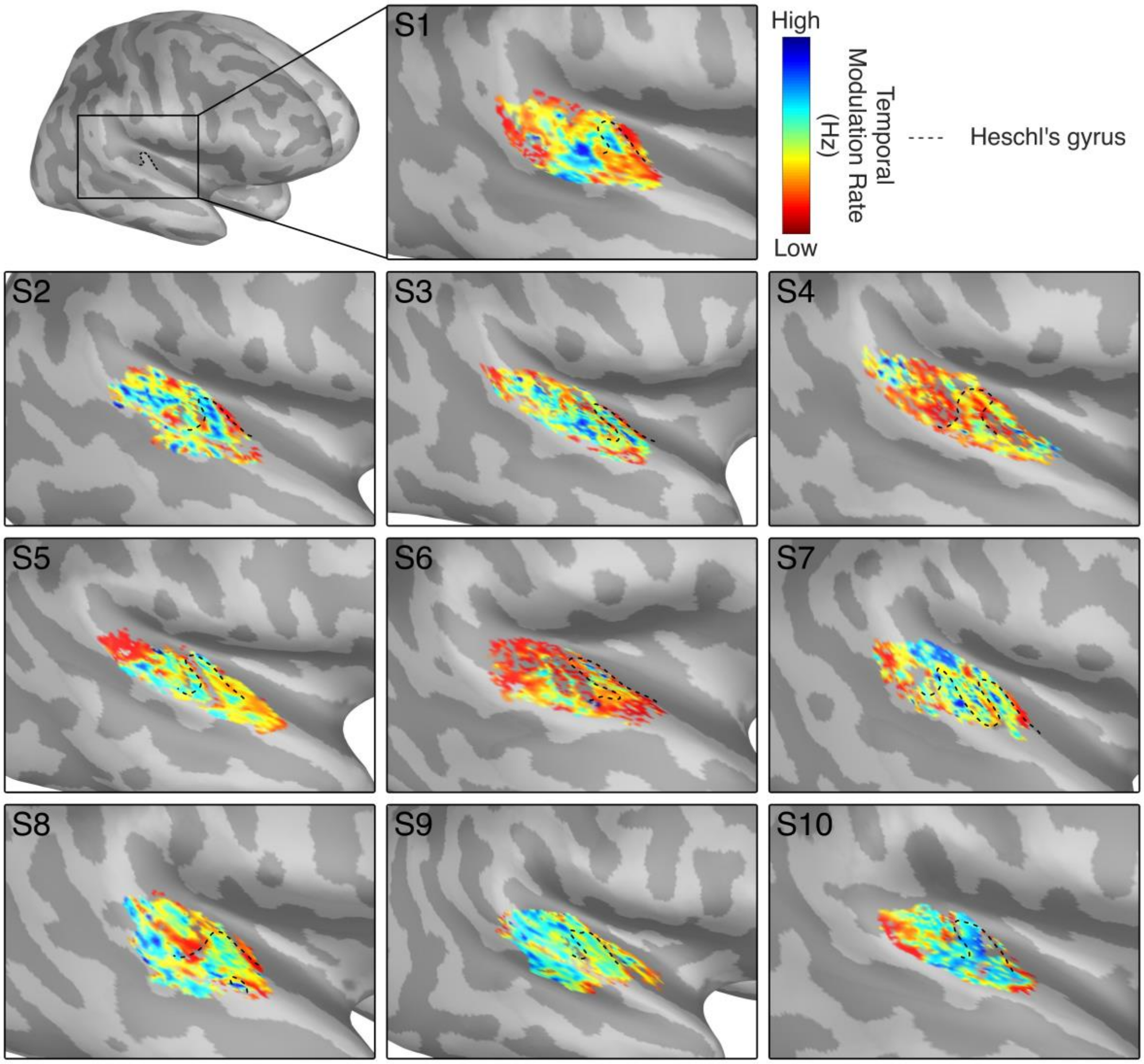


***Figure S3. Right Hemisphere Temporal Modulation Rate Maps.*** *Temporal modulation rate maps are shown for subjects 1 to 10. Heschl's gyrus is outlined for reference. Colormap limits are set near the local minima and maxima of each subject to best visualize gradient patterns.*

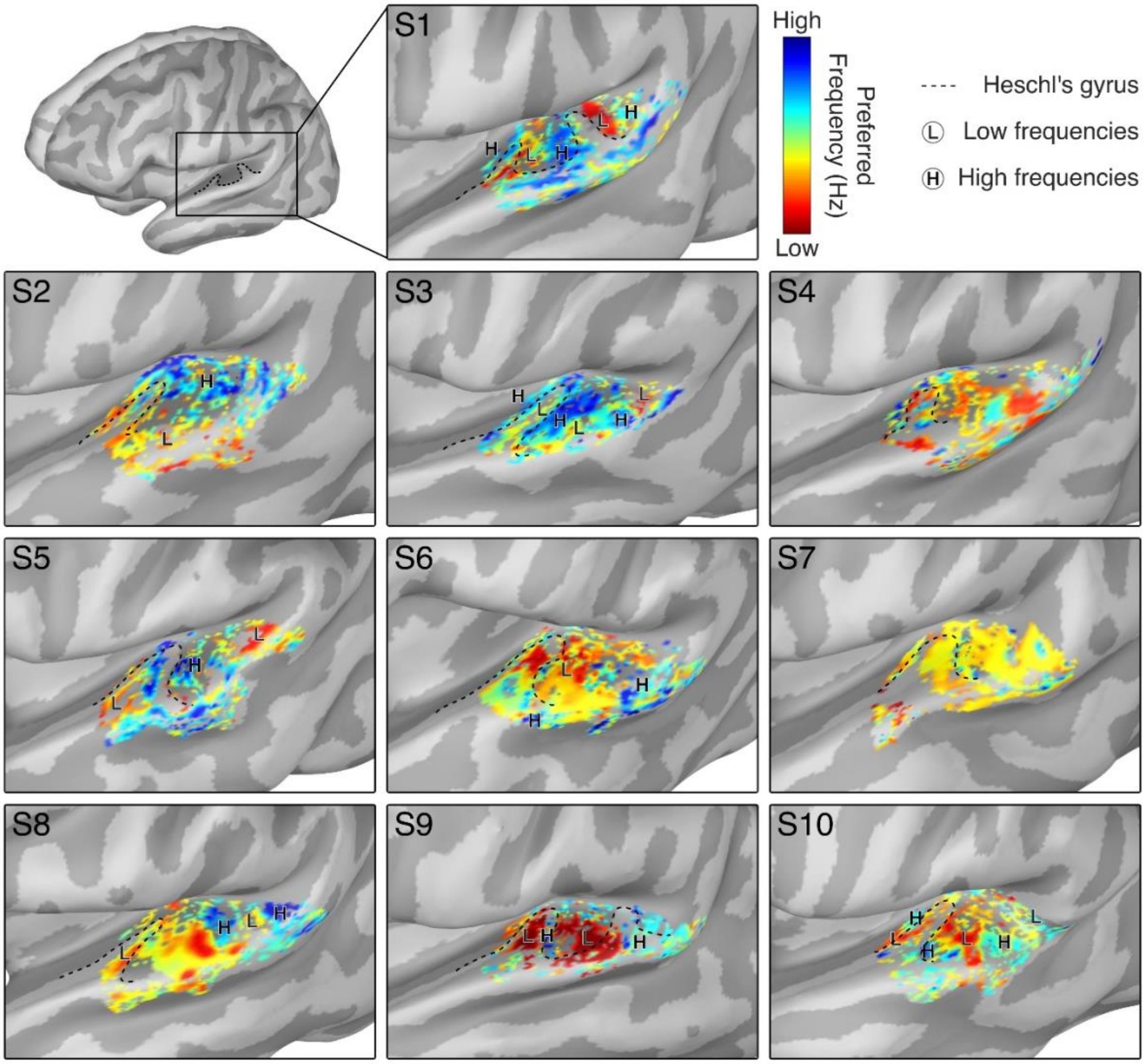


***Figure S4. Left Hemisphere Best Frequency Maps.*** *Best frequency maps are shown for subjects 1 to 10. Major regions of high and low frequencies are marked as H and L, respectively, and Heschl's gyrus is outlined for reference. Clear tonotopic gradients could not be identified for S4 and S7. Colormap limits are set near the local minima and maxima of each subject to best visualize gradient patterns.*

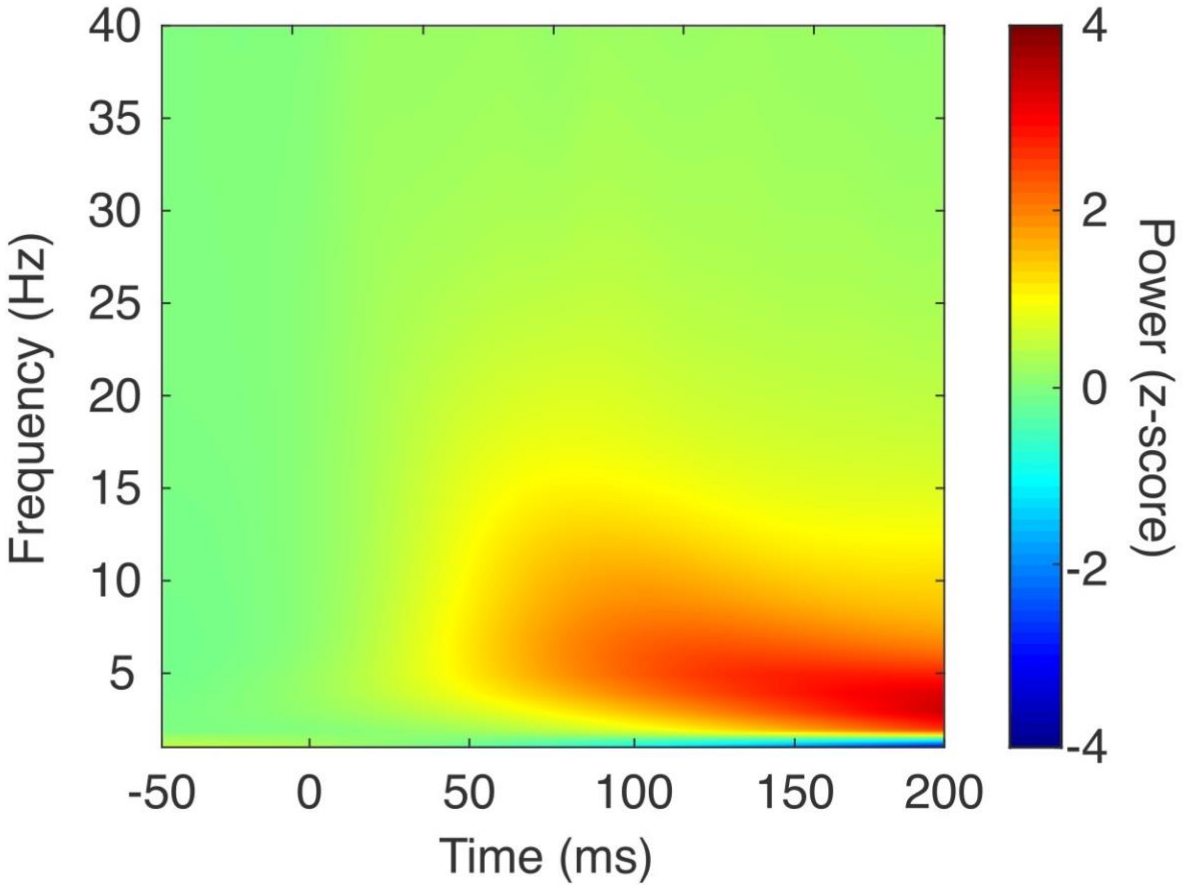


***Figure S5.*** *Average time-frequency decomposition of the isointensity pure tones stimulus train. Morlet wavelet time-frequency decomposition of the normalized power of each frequency over Heschl gyrus during a train of isointense pure tones. The decomposition shown here is the average across all-subjects. By applying a threshold of 1 on the z-score normalized power, most of the frequencies of interest at the 100 ms latency (the M100 response) can be isolated to the frequency band spanning 3-13 Hz, starting around 100 ms after the isointensity pure tone stimulus.*

| Subject | *Lower cutoff frequency (Hz)* | *Upper cutoff frequency (Hz)* |
|---|---|---|
| S1 | 3 | 8 |
| S2 | 4 | 10 |
| S3 | 4 | 13 |
| S4 | 3 | 10 |
| S5 | 4 | 10 |
| S6 | 4 | 12 |
| S7 | 4 | 8 |
| S8 | 3 | 11 |
| S9 | 4 | 8 |
| S10 | 3 | 8 |

***Table S1. Optimal lower and upper frequency cutoffs for the band-pass filter of each subject.*** *The optimal lower and upper frequency cutoffs are shown for each subject, as obtained through the time frequency analysis described in the Methods section. From these cutoff numbers, a minimum lower cutoff frequency of 3 Hz and a maximum cutoff frequency of 13 Hz can be obtained. These cutoffs are used to define the 3-13 Hz bandpass filter applied to the isointensity pure tone stimulus used in our analysis.*
